# Metabolic Adaptations of Prostate Cancer Cells Under Chronic Fructose Stimulation

**DOI:** 10.1101/2025.07.07.663510

**Authors:** Carolina E. Echeverría, Vanessa I. Oyarzún, Claudia Jara, Emilia Flores-Faúndez, Catalina Ramírez, Verónica Torres-Estay, Cheril Tapia-Rojas, Jorge Díaz, Manuel Meneses, Jorge Cancino, Alejandro Pinilla, Alejandro San Martín, Spencer Rosario, Alejandro S. Godoy

**Author notes:** Corresponding author: Alejandro S. Godoy.

## Abstract

**Background:** Dietary sugars, particularly fructose, are increasingly implicated in cancer progression through their impact on tumor metabolism. However, the specific metabolic adaptations driven by fructose in prostate cancer (PCa) remain unexplored.

**Methods:** We investigated the metabolic consequences of chronic fructose exposure in androgen-sensitive (LNCaP) and androgen-independent (PC-3) PCa cell lines. We evaluated the expression, localization, and function of the fructose transporter Glut-5 and assessed metabolic fluxes, enzyme expression, lipid accumulation, and global metabolite profiles using molecular, imaging, and metabolomic approaches.

**Results:** Glut-5 was primarily localized to early endosomes under basal and fructose-stimulated conditions, suggesting a non-canonical role potentially consistent with transceptor function. Chronic fructose exposure significantly upregulated Glut-5 expression and enhanced fructose uptake, but did not alter substantially its subcellular localization. Functionally, fructose reduced lactate production and mitochondrial ATP output, indicating a metabolic shift away from glycolysis and oxidative phosphorylation. In LNCaP cells, fructose induced robust activation of de novo lipogenesis (DNL), evidenced by upregulation of FASN and G6PD, increased lipid droplet accumulation, and enhanced levels of key fatty acid metabolites (e.g., TG, EPA, DHA). In contrast, PC-3 cells exhibited a distinct metabolic response, characterized by increased ceramide and amino acid metabolites. Notably, pharmacological inhibition of lipid metabolism using etomoxir abrogated proliferation in both cell lines under fructose stimulation.

**Conclusions:** These findings reveal that fructose promotes a metabolic reprogramming in PCa cells that is cell type- and AR-dependent, enhancing lipogenesis and potentially contributing to tumor progression. Our study identifies Glut-5–mediated fructolysis and lipid metabolic pathways as key vulnerabilities in PCa, offering potential avenues for metabolic intervention.

## Introduction

Prostate Cancer (PCa) is the most prevalent malignancy among men over the age of 50 in countries with high and very high human development indices^1^. In the United States of America alone, projections estimated 313,780 newly diagnosed cases and 35,770 deaths attributed to this disease by 2025^2^. This significant incidence of PCa correlates with a defining characteristic of developed nations: the modern lifestyle characterized by lack of physical activity and excessive calorie intake, particularly from sugars^3^.

Among the most consumed sugars, fructose plays a significant role in contributing to various metabolic-associated diseases. Excessive consumption of fructose has been linked to an increase in visceral adiposity^4^, insulin resistance^5^, and cancer^6–9^. Consequently, numerous research groups have turned their focus toward exploring the correlation between fructose consumption and various types of cancer^10–12^. Our group has developed a specific interest in unraveling the complex relationship between fructose consumption and PCa^13,14^. Carreño *et al*^11^., reported compelling findings indicating that chronic fructose stimulation leads to increased proliferation of PCa cells and promotes tumor growth. These results were observed through both *in vitro* and *in vivo* models, shedding light on the potential mechanisms by which fructose may influence the progression of PCa. Even more intriguingly, Carreño *et al*^11^., discovered that individuals with PCa displayed notably elevated levels of fructose in their serum compared to those with benign conditions. Similar findings have been observed in patients diagnosed with pancreatic cancer, further highlighting the potential systemic impact of chronic fructose exposure^15^. Beyond the biological implications of this finding, these results suggest that PCa cells are likely exposed to sustained high concentrations of fructose via the bloodstream, which may influence their metabolic and proliferative behavior.

PCa cells have the capacity to invade nearby seminal vesicles (SVs). SVs are known for having one of the highest levels of fructose in the human body^16^. This sugar serves as a critical metabolic substrate for sperm, enabling them to generate ATP and support cell motility^14^. PCa cells may also gain access to fructose via the seminal fluid secreted by the SVs, which may be enhanced by local invasion of the PCa cells into the SVs, a process called seminal vesicle invasion (SVI). Interestingly, SVI is an adverse pathologic factor conferring a high rate of PCa recurrence over and above other local invasion^17^. Therefore, it is tempting to speculate that SVI is especially hazardous since it provides access to fructose as an alternative energy source for PCa cells to utilize upon its invasion to SVs and that this fructose-enriched environment might boost the aggressiveness of PCa cells resulting in a worse PCa patient outcome. In support of this hypothesis, previously published results from our laboratory indicate that chronic fructose exposure promote the invasion capability of the LNCaP and PC-3 PCa cell lines cultured *in vitro*^11^. Fructose is transported into human cells primarily through facilitative hexose transporters, known as Gluts^13,18^. Among these, the isoform Glut-5 is uniquely significant as the most highly overexpressed fructose transporter in malignant cells^6,12,19,20^. Glut-5 is distinguished from other Glut isoforms by its exclusive specificity for fructose and is predominantly expressed on the apical surface of intestinal epithelial cells, as well as in the kidney, liver, and testis^13^. Prior studies have demonstrated substantial upregulation of Glut-5 in PCa tissues relative to benign prostate tissues^10,11^. Moreover, *in vitro* kinetic analyses have reported a Michaelis-Menten constant (*K_m_*) of 6.8 mM in LNCaP cells and 7.1 mM in PC-3 cells, underscoring Glut-5 as the primary mediator of fructose transport in these malignant cells. Interestingly, immunohistochemistry analyses of PCa cells and other tumor models have revealed a substantial proportion of Glut-5 immunostaining localized intracellularly^17–19^. However, the subcellular localization and trafficking dynamics of Glut-5 under both physiological and pathological conditions remain poorly understood.

Fructose metabolism is intricately associated with the aggressiveness and progression of various cancers, including breast^21^, colorectal cancer^6^, lung^22^, and prostate^11^. Similar to glucose, fructose serves as a substrate for several metabolic pathways, such as the synthesis of fatty acids through the *de novo* lipogenesis (DNL) pathway and nucleotides via the oxidative and non-oxidative pentose phosphate pathway (PPP), both of which are critical for cellular replication and tumor growth. In pancreatic cancer, fructose has been shown to enhance nucleotide synthesis via the non-oxidative PPP, mediated by the upregulation of transketolase (TKT)^23^. In contrast, studies on colorectal cancer models have demonstrated that fructose stimulates the DNL pathway through the upregulation of fatty acid synthase (FASN)^6^. These findings collectively suggest that the molecular mechanisms through which fructose promotes malignant cell proliferation and tumor progression are highly tumor-specific. Nevertheless, the metabolic pathways or fundamental metabolic profiles altered by chronic fructose exposure in PCa cells remain largely undefined, necessitating further exploration.

This study seeks to investigate the metabolic adaptations of PCa cells in response to prolonged fructose exposure, with a focus on the subcellular localization and redistribution of Glut-5, a critical fructose transporter, following fructose stimulation. Additionally, this research examines the metabolic fluxes involved in fructose metabolism using *in vitro* models. By elucidating the metabolic alterations induced by chronic fructose exposure, this study aims to enhance our understanding of the mechanisms driving PCa progression and identify potential metabolic vulnerabilities that could serve as therapeutic targets.

## Materials and Methods

### Cell culture

All human prostate cancer cell lines, LNCaP (androgen sensitive) and PC-3 (metastatic), were commercially obtained from the ATCC (American Type Culture Collection) and used for no more than 30 passages. Cells were maintained at 37°C in a humidified atmosphere of 5% CO_2_ and cultured in RPMI-1640 Medium (Gibco-Life Technologies) supplemented with 10% FBS (Fetal Bovine Serum) (Gibco-Life Technologies), 1% penicillin/streptomycin (Life Technologies, Carlsbad, US) and 1nM of DHT. All cell lines were tested every 2 weeks for the presence of *Mycoplasma*. Sugar-free RPMI (Gibco-Life Technologies) was utilized in hexose stimulation. Glucose (#G7021-100G, Sigma) and fructose (#F0127-100G, Sigma) powders were dissolved in water to prepare a 1 M stock solution, which was subsequently diluted in sugar-free RPMI to obtain working solutions.

For hexose stimulation, LNCaP and PC-3 cells were cultured in sugar-free RMPI containing 5.5 mM glucose or 5.5 mM fructose, 10% FBS and 1% penicillin/streptomycin. Cells were maintained under glucose or fructose stimulation for short periods of stimulation (24, 48, 72 hours) or long periods of stimulation (10 days), replacing the media every 48 hrs.

### Cell viability assays

For cell viability assay, cells were seeded on a 12-well culture plate and cultured to 90% confluency at the time of the readout. The day after seeding, cells were treated with different fructose concentrations (0.01 mM, 0.5 mM or 5.5 mM) and incubated for 24 h, 48 h, and 96 h. To asses cell viability, relative cells numbers were measured using an automated counter (LUNA, Logos Biosystems) according to the manufactures protocol.

### Western blot

Cultured cells were pelleted and lysed in RIPA lysis buffer [50 mmol/L Tris (pH 8), 1% Triton x-100, 0.1% SDS, 150 mmol/L NaCl, 5 mmol/L EDTA, and 0.5% sodium deoxycholate] supplemented with 1X protease inhibitor cocktail (#11836170001-Roche) and separated by centrifugation. Protein extract (determined by BCA method (#23227-Thermo Scientific)) was subjected to electrophoresis in a 10-12% SDS-PAGE and then transferred onto a 0.45 µm PVDF membrane. Then, PVDF membranes were blocked with 5% (w/v) BSA (Bovine Serum Albumin) in 1X TBS-Tween (Tris-buffered saline with 0.1% (v/v) Tween 20) (TBST) for 1 hour at room temperature. Then, PVDF membranes were incubated overnight with primary antibodies, such as anti-Glut-1 (#GT12-A), anti-Glut-2 (#GT22-A), anti-Glut-5 (#GT52-A), anti-Glut-7 (#GT73A), anti-Glut-9 (#GT91-A), anti-β-actin (#A1978), anti-G6PD (#SC-373886), anti-TKT (#SC-390179), anti-FASN (#SC-48357), anti-vinculin (#ab18058), anti-LDH-A (#CST2012S) and anti-p-LDH-A (#CST8176S) diluted in 5% BSA at 4°C. After an overnight incubation, a TBST wash, and the appropriate HRP-conjugated secondary antibodies (Agilent Dako) were incubated for 1 hour at room temperature. Lastly, the reaction was revealed by the ECL Western Blot Analysis System kit (#34075, Thermo Scietific), using a G Box system (Syngene).

### RNA extraction, reverse transcription, and quantitative real-time PCR

Total RNA from LNCaP and PC-3 cells was isolated directly from plates using Trizol® (Invitrogen). For quantitative real-time PCR (qRT-PCR), 1.0 µg RNA was reversed-transcribed using M-MLV reverse transcriptase set (Promega, Madison). The resulting cDNA was diluted 1:5 and qPCR were performed with Fast SYBR Green Mastermix (Applied Biosystem) in the StepOne Real-Time PCR System (Applied Biosystem). The relative expression of each gene was calculated by comparative △C_t_ method after normalizing to endogenous control (18S). Primer sequences for qRT-PCR are detailed in the supplementary table I.

### Lactate production (ELISA kit)

To assess the changes in lactate levels of PCa cells during fructose stimulation, LNCaP and PC-3 cells were stimulated with 5.5 mM glucose (as control) or 5.5 mM fructose for 48h and 10 days. Cellular pellets and supernatants were collected for measurement of lactate concentration. Intracellular and extracellular lactate was measured by using a Lactate assay kit (#ab65330 Abcam) according to the manufacture’s protocol for colorimetric assay. All samples were deproteinized with the Deproteinizing Sample Preparation Kit-TCA (#ab204708 Abcam) before analysis. Lactate concentration was determined using a standard curve and adjusted to the relevant dilution factor.

### Lactate production (Forster Resonance Energy Transfer [FRET]-based lactate sensor)

For acute intracellular lactate production measurements, cells were incubated with 2 mM glutamine for 48 hours and during the experiment’s cells were exposed to 5 mM glucose or 5 mM fructose. For laconic gene delivery and expression, LNCaP and PC3 cells were transfected at 60% confluence using Lipofectamine 3000 (Gibco) and then infected with 5×10^6^ pfu adenoviral particles of Ad-Laconic (Vector BioLabs). After 24 hours of expression covers slides were mounted in a superfusion chamber. All experiments were performed at room temperature in a superfusion solution of the following composition: 136 mM NaCl, 1.25 mM CaCl_2_, 1.25 mM MgSO_4_, 10 mM HEPES, 5 mM KCl, pH 7.4. Cells were imaged with an upright Olympus FV1000 confocal microscope equipped with a 20 water-immersion objective (numerical aperture, 1.0) and a 440nm solid-state laser. Laconic (Addgene plasmid #44238)^24^, was excited at 430 nm for 0.2– 0.8s and was detected at 485/40 nm [cyan fluorescent protein or monomeric teal fluorescent protein (mTFP)] and 535/30nm (Venus). Time series images were taken every 10s in XYT scan mode (scan speed: 156 Hz; 800 × 800-pixel; pinhole 800 μm). The ROI (region of interest) was selected from a cytosolic homogeneous single-cell signal. The background was subtracted from each emission channel and the resulting mTFP/Venus ratio was used to calibrate the data. Two-point calibration protocol was utilized to covert the FRET signal into lactate concentrations. RMIN was achieved with 6 mM oxamate and RMAX was obtained from the maximal signal in presence of AR-C155858 for each cell and experiment. The K_d_ values were obtained from original Laconic paper^24^. The intracellular lactate concentrations were obtained interpolating the mTFP/Venus ratio values into a doble-rectangular hyperbole curve.

### Cellular ATP content

Total cellular ATP levels were measured in PC-3 and LNCaP cell lysates obtained with HEPES buffer (25mM HEPES, 125 mM NaCl, 25mM NaF, 1 mM EDTA, 1mM EGTA, 1% NP-40, pH = 7,4) using a luciferin/luciferase bioluminescence assay kit (ATP Determination Kit #A22066, Molecular Probes, Invitrogen). The amount of ATP in each sample was calculated from standard curves and normalized to the total protein concentration.

### ManCou1-H Microplate Uptake

For microplate assay, LNCaP and PC-3 cells were cultured for 10 days under fructose or glucose treatment. On day 7, cells at ∼80% confluence were harvested and seeded into black 96-well flat-bottom plates (20,000 cells/well), then allowed to grow for 3 more days to complete a total of 10 days of treatment. Cells were subsequently washed with pre-warmed (37°C) PBS solution, treated with ManCou1-H probes diluted in PBS, and incubated at 37°C and 5% CO2 for 10 minutes. After incubation, cells were carefully washed with warm PBS (3 × 100 µL), and endpoint fluorescence was immediately measured using a Synergy HTX Multimode plate reader controlled by the Gen5 software (version 3.12) with excitation at 355nm, emission at 460 nm, and a read time of 1.0 s). Uptake studies were conducted using black 96 - well flat-bottom plates. Fluorescence of Mancou1-H probes in cells was measured in the presence of 5.5 µM fructose or glucose. For this purpose, PBS solution containing 20 µM ManCou1-H and the indicated sugar concentration was prepared and added to the wells.

### Isolation of a mitochondrial-enriched fraction

A fraction enriched in mitochondria was isolated from PC-3 and LNCaP cell lysates obtained with MSH buffer (230 mM mannitol, 70 mM sucrose, 5 mM Hepes, pH 7.4) supplemented with phosphatase (NaF 20x, Na2P2O7 300x, Na3VO4 100x) and protease (#78429 Thermo Fisher Scientific) inhibitors. Lysates were centrifuged at 600 g for 10 min at 4 °C. The pellet was discarded, and the supernatant was centrifuged at 8000 g for 10 min at 4 °C. The resulting pellet was resuspended in KCl respiration buffer (125 mM KCl, 0,1% BSA, 20 mM HEPES, 2 mM MgCl2, 2.5 mM KH0PO4, pH = 7.2).

### Mitochondrial ATP Production

For measurement of mitochondrial ATP production, 25 µg of mitochondria protein were incubated with oxidative substrates in KCl respiration buffer for 30 min at 37°C, then were centrifuged at 8000 g for 10 minutes at 4°C, and the ATP concentration was measured in the supernatant using the luciferin/luciferase bioluminescence assay kit (#A22066 Invitrogen). The amount of ATP in each sample was calculated from standard curves and normalized to the total protein concentration.

### Mitochondrial O_2_ consumption

For measurement of O_2_ consumption, 25 µg of mitochondria protein were incubated with oxidative substrates and the O_2_ reagent (Extracellular Oxygen Consumption Reagent, #ab197242 Abcam) in KCl respiration buffer for 30 min at 37°C. The extracellular oxygen consumption reagent dye is used with an oil layer which is added on top of the assay medium to limit diffusion of oxygen into the assay medium. As mitochondrial respiration depletes the oxygen within the assay medium, quenching of the fluorescent dye is reduced, and the fluorescence signal increases proportionately.

### Lipid staining assay

Lipids were stained using oil red O (ORO). PCa cells under chronic treatments were washed with PBS and fixed with 4% paraformaldehyde (PFA) for 15 min at room temperature (RT). Prior to staining, a stock solution of 3 mg/ml was prepared in isopropanol. The stock solution was diluted in water (3:2) and filtered (0.22um) prior to staining the cells. Fixed cells were washed with PBS, and then incubated with 60% isopropanol for 5 min. After this, the filtered ORO working solution was added for 15 min and then washed with PBS. Dapi was added for 10 min at RT to visualize nuclei. Cells were washed with PBS and mounted onto glass slides using Prolong Diamond Antifade (Molecular Probes) and allowed to cure for 24 hours. Images were acquired under SP8 Confocal Microscope. At least three randomly selected microscopic fields were taken for each condition.

### Immunohistochemistry

Antigen retrieval was performed using 0.01 mol/L sodium citrate buffer pH 6 for 30 minutes^11,19^. Endogenous peroxidase activity was inhibited by using 3% H_2_O_2_ in methanol and nonspecific binding of the antibody was blocked with 2% BSA for 20 minutes. Tissue sections were incubated with anti-Glut-5 (1:100, Alpha Diagnostic International) antibody for 12 hours. Tissue sections were then incubated with horseradish peroxidase (HRP)-conjugated secondary antibody (1:100 Dako) for 1 hour at room temperature. Peroxidase activity was developed using 3,3-Diaminobenzidine (Dako) and H_2_O_2_. Harrýs hematoxylin was used to counterstain nuclei. Slides were dehydrated through graded alcohol to xylene and mounted with coverslips using Prolong Diamond Antifade (Molecular Probes).

### Immunofluorescence

Cells were seeded on a glass coverslip (12 mm) into a 24-well plate in RPMI1640 medium supplemented with 10% FBS. After 24 hours, cells were incubated with glucose-free RPMI1640 medium supplemented with 5.5 mM glucose or 5.5 mM fructose for 10 days and cells were fixed with 4% PFA for 15 minutes at RT. Cell permeabilization was performed using 0.1% Triton X-100 in 1X PBS/Ca^+^/Mg^+^ for another 15 minutes. Subsequently, cells were incubated for 1 hour at 37°C with primary antibodies anti-GM130 (bd-610822), anti-NaK (sc-514614), anti-Calnexin (sc-23954), anti-transferrin (T233660), anti-EEA1 (sc-365652), anti-TKT (sc-390179), anti-FASN (sc-48357), anti-G6PD (sc-373886), anti-Glut-1 (GT12-A), anti-Glut-5 (GT52-a) and anti-Glut-5/blocking peptide (GT52-P). After washing with PBS, cells were nuclear counterstained with Dapi for 10 minutes at 37°C. Cells were washed with PBS and mounted onto glass slides using Prolong Diamond Antifade (Molecular Probes) and allowed to cure for 24 hours. At least three randomly selected microscopic fields were taken for each condition. Images of fluorescent-stained cells were acquired on SP8 Confocal Microscope. Raw.tif files were processed using FIJI (Image J) and/or Canvas Draw (Canvas GFX) to create stacks, adjust levels, and/or apply color.

### Effect of Etomoxir on fructose-stimulated PCa cell proliferation using the Operetta cell analyzer

Cells were seeded on a glass coverslip (12 mm) into a 24-well plate in RPMI1640 medium supplemented with 10% FBS. After 24 hours, cells were incubated with glucose-free RPMI1640 medium supplemented with 5.5 mM glucose or 5.5 mM fructose for 10 days and cells were stimulated with or without Etomoxir for 0 to 72 h.

### Biocreates pepline

Samples were prepared and analyzed in the Roswell Park Comprehensive Cancer Center Bioanalytics, using the MxP Quant 500 kit (Biocrates Life Sciences AG) in accordance with the user manual. Cell pellet samples were resuspended in a ratio of 25 μL of solver (85% ethanol and 15% 0.01 M phosphate buffer) to 1×10^6^ cells. The resuspended cells went through cycles of sonication (in ice-bath) and snap freezing on dry ice. Samples were then centrifuged to obtain a supernatant for analysis. A 10 μL aliquot of each cell pellet supernatant was added on the filterspot (already containing internal standard) in the appropriate wells. The plate was then dried under a gentle stream of nitrogen. The samples were derivatized with phenyl isothiocyanate (PITC) for the amino acids and biogenic amines, and dried again. Sample extract elution was performed with 5 mM ammonium acetate in methanol. These extracts were diluted with either water for HPLC-MS/MS analysis (1:1) or kit running solvent for low injection analysis (FIA)/MS/MS (50:1) using Shimadzu HPLC system interfaced with a Sciex 5500 mass spectrometer. Data was processed using MetIDQ software, and Limma for differential metabolite analysis.

### Statistical analysis

All statistical data analysis was performed using GraphPad Prism 9.0 through unpaired Student *t* test, one-way ANOVA or Mander’s test as appropriate. *P* < 0.05 was set as a threshold for significant results.

## Results

### Glut-5 is expressed and localized to early endosomal compartments in prostate cancer cells

Initially, we verified the specificity of the anti-Glut-5 antibody by pre-incubating it with increasing concentrations (100 and 200 µg/mL) of the anti-Glut-5 blocking peptide before conducting the immunofluorescence (IF) procedure in LNCaP and PC-3 cell lines (Suppl. Fig. 1). This approach is prudent because many commercial antibodies are raised against peptide sequences from the C- or N-terminus, which may have epitope similarity across multiple isoforms of solute transporters. As depicted in Supplementary Fig. 1A-B, Glut-5 staining intensity significantly decreased in a concentration-dependent manner of the blocking peptide in both cell types. To confirm the specificity of the anti-Glut-5 blocking peptide, we pre-incubated the highest concentration of the blocking peptide (200 µg/mL) with the anti-Glut-1 antibody under the same experimental conditions. As anticipated, no decrease in Glut-1 staining intensity was observed when the anti-Glut-1 antibody was pre-incubated with the anti-Glut-5 blocking peptide (Supp. Fig. 1C-D). This results ultimately confirmed the specificity of the Glut-5 antibody.

We examined the expression of Glut-5 mRNA and protein in LNCaP and PC-3 cell lines under basal conditions (cell cultured in regular media). Glut-1 mRNA and protein expression were used as controls (Fig. 1A-C). Our results indicated that Glut-1 mRNA was significantly overexpressed in PC-3 cell line when compared to LNCaP cell line (Fig. 1A, Glut-1 mRNA). However, no significant differences were observed in Glut-5 mRNA expression between the two cell lines (Fig. 1A, Glut-5 mRNA). The correlation between mRNA levels and protein levels for solute transporter proteins is not always strong due to multiple layers of post-transcriptional, translational, and post-translational regulation. Therefore, we analyzed the Glut-1 and Glut-5 proteins expression in LNCaP and PC-3 cell lines (Fig. 1B-C). Contrary to our mRNA level findings, the IF results indicated no significant differences in Glut-1 and Glut-5 protein levels between LNCaP and PC-3 cell lines (Fig. 1C). Glut-1 protein tended to decrease its expression in PC-3 as compared to the less aggressive LNCaP cell line, and Glut-5 protein exhibited the opposite behavior.

**Figure 1.**
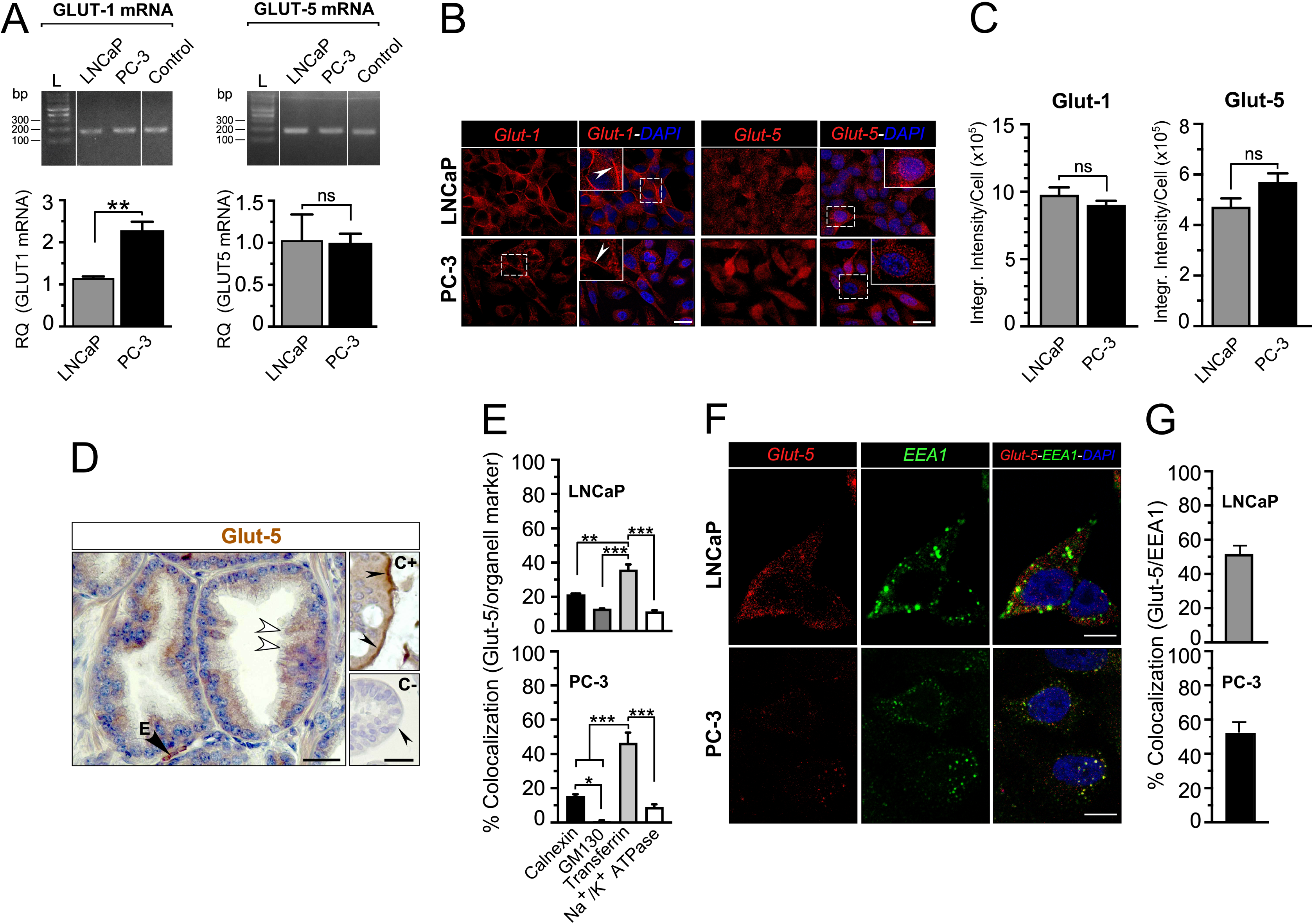
Glut-5 is localized in endosomal compartments in prostate cancer cells. (A) PCR analysis on an agarose gel using specific primers for Glut-1 and Glut-5 in LNCaP and PC-3 cells under basal conditions (top panels). Human duodenum was used as a positive control. 18S was used for data normalization and gene expression analysis in LNCaP and PC-3 cells (n=3). t-test: ns (not significant), **p<0.01. (B) Immunofluorescence staining of Glut-1 and Glut-5 in LNCaP and PC-3 cells under basal conditions. DAPI was used to counterstain nuclei. Glut-1 shows a membrane localization pattern in both cell types (Insets, white arrowheads), whereas Glut-5 is primarily localized intracellularly (Insets). Scale bar: 10 µM. (C) Quantification of glucose transporter expression under basal conditions, measured as integrated intensity normalized per cell (n=3; t-test: ns, not significant). (D) Immunohistochemistry of Glut-5 in prostate cancer tissue (Gleason grade 6). E: erythrocyte. Human small intestine tissue was used as a positive control. For the negative control, 2% BSA replaced the primary antibody. Scale bar: 20 µm. (E) Quantification of Glut-5 colocalization with each organelle marker in LNCaP and PC-3 cells under basal conditions (n=3; *p<0.05, **p<0.01, ***p<0.0001; one-way ANOVA, Manders coefficient). (F) Immunofluorescence staining of Glut-5 and the early endosome marker (EEA1) in LNCaP and PC-3 cells under basal conditions. Scale bar: 5 µM. (G) Quantification of Glut-5 colocalization with EEA1 in PCa cell lines under basal conditions.

We next examined the localization of the transporter proteins. Consistent with expectations for a transport protein, Glut-1 protein showed a pattern of expression that suggests primarily a plasma membrane localization (Fig. 1B, Glut-1 and Glut-1-DAPI [inset, arrow head]) in both cell lines. Remarkably, Glut-5 was mostly expressed in the cytoplasm with a punctate pattern homogeneously distributed throughout the cytoplasm (Fig. 1B, Glut-5 and Glut-5-DAPI [inset]), suggesting an endosomal compartment localization. To validate this finding, we examined Glut-5 expression in ten human PCa tissue specimens. Eight out of ten specimens displayed varying intensities of Glut-5 immunostaining, with a pattern suggesting intracellular localization of the transporter (Fig. 1D, white arrowheads). The specificity of the anti-Glut-5 antibody was validated by two ways: 1) by detecting the Glut-5 protein at the plasma membrane of erythrocytes, as observed in clinical specimens of PCa (Fig. 1D, black arrow, E) and 2) by detecting the Glut-5 protein at the apical membrane of enterocytes in human intestinal tissue (Fig. 1D, inset C+, black arrow heads). In the latter case, the absence of the primary antibody served as a negative control (Fig. 1D, inset C-, black arrow head).

Previous studies from our^10,11,19^ and others^25,26^ have consistently reported that Glut-5 protein in malignant cells from human tissue specimens of different origins was mainly localized to the cytoplasm. However, very few studies^27^ have explored the subcellular localization of Glut-5 in cancer cells, both *in vitro* and *ex vivo*. In this study, we undertook a comprehensive analysis of Glut-5 localization in LNCaP and PC-3 cell lines utilizing laser confocal scanning microscopy (Fig. 1E-H). We performed immunofluorescence assays employing various organelle-specific markers, including an endoplasmic reticulum (ER) marker (Calnexin) and a Golgi apparatus (GA) marker (GM130). Furthermore, our analysis encompassed markers for the endosomal compartment (transferrin and EEA1) and the plasma membrane (Na^+^/K^+^ ATPase) (Fig. 1E-G). Notably, we observed that Glut-5 predominantly colocalized with transferrin-staining intracellular compartments in both cell types (Fig. 1E-F). Semi-quantitative analyses revealed that approximately 40% and 55% of Glut-5 immunostaining colocalized with transferrin-staining compartments in LNCaP and PC-3 cells, respectively (Fig. 1F). A lower percentage of Glut-5 colocalization was observed with the ER (21% for LNCaP and 15% for PC-3) and GA (12% for LNCaP and 0.98% for PC-3) markers (Fig. 1D). Remarkably, in both LNCaP and PC-3 cell lines, 5% or less of the Glut-5 immunostaining was localized to the plasma membrane (Fig. 1F). These data suggest that Glut-5 is primarily localized within early endosomal compartments.

Transferrin molecules are continually recycled from the cell membrane to early endosomes^28^. Consequently, we performed immunofluorescence staining for early endosome compartments using the EEA1 marker. Notably, we observed significant colocalization of EEA1 and Glut-5 in both cell types (Fig. 1G). Semi-quantitative analysis revealed approximately 55% and 60% colocalization of Glut-5 with EEA1 in LNCaP and PC-3 cells, respectively, under basal conditions (Fig. 1G). Collectively, these data suggest that, at steady state, Glut-5 is predominantly localized to the early endosomal compartment in both LNCaP and PC-3 cells.

### Chronic fructose stimulation upregulates the expression of Glut-5 without altering its subcellular localization

Previous studies by Liang *et al*.^12^ found that PC-3 cells cultured chronically in the presence of fructose increased their expression of Glut-5 and developed fructolysis *in vitro*. First, we validated that chronic exposure to glucose or fructose, as the sole sugar in the culture medium, does not affect tumor cell survival. Our results showed that both glucose and fructose sustained over 95% survival in LNCaP and PC-3 cells across all tested time points, from 0 to 96 hours (Supplementary Fig. 2). This analysis was also repeated after 7 and 10 days of incubation, yielding similar results (data not shown).

Next, we began examining the expression levels, functionality, and, more importantly, the subcellular localization of Glut-5 in response to chronic fructose stimulation in LNCaP and PC-3 cells. Analyses of Glut-1 were included in this study as an internal control (Fig. 2). LNCaP and PC-3 cells were cultured with sugar-free RMPI medium containing either 5.5 mM fructose or 5.5 mM glucose for 10 days, which mimics physiological hexose concentrations^11^. We found that the level of *SLC2A5* gene expression was elevated after fructose treatment in either LNCaP (1.5-fold change) or PC-3 (1.6-fold change) cells when compared to control condition (Fig. 2A). Although the results were not statistically significant, the expression of the *SLC2A1* gene exhibited a tendency towards downregulation in the presence of fructose across both cell lines. Consistent with the mRNA expression analyses, elevated Glut-5 protein expression was observed in the whole cell lysates by approximately 4.0 folds for LNCaP cells and 2.2 folds for PC-3 cells, after incubation with fructose for 10 days (Fig. 2B-C). Consistently with the qPCR analyses, Glut-1 protein expression was not significantly affected by the presence of fructose (Fig. 2B-C). In addition, we examined the expression of the other members of the Glut family with the potential to transport fructose, which included Glut-2, Glut-7, Glut-9, and Glut-11^13^ (Fig. 2B). Chronic fructose stimulation did not affect the expression of any of these transporters.

**Figure 2.**
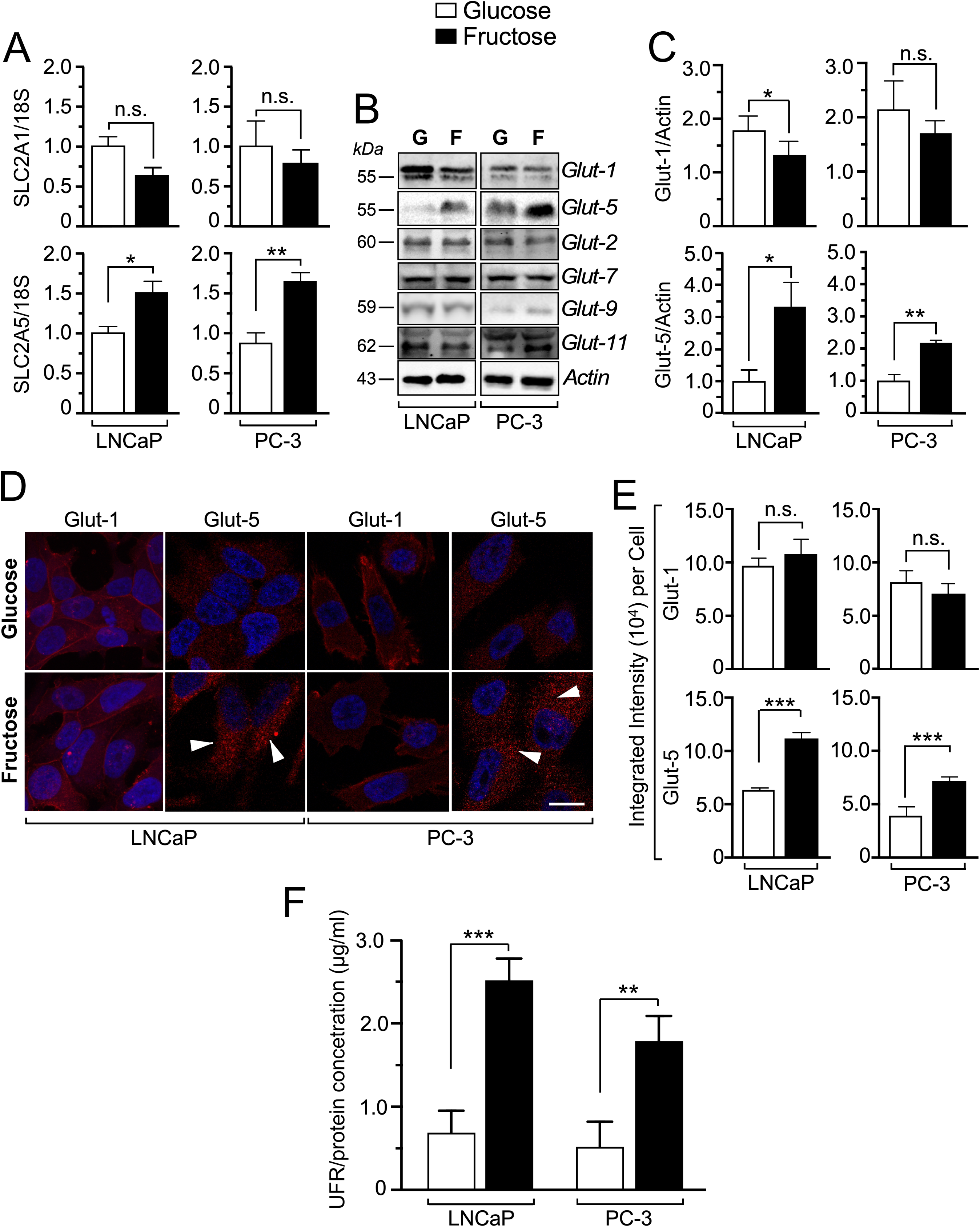
Chronic fructose stimulation increases *SLC2A5*/Glut-5 expression in prostate cancer cell lines *in vitro*. (A) mRNA levels of *SLC2A1* and *SLC2A5* in LNCaP and PC-3 cells following 10 days of stimulation with 5.5 mM glucose (white bars) or 5.5 mM fructose (black bars) (n=3–4; ns: not significant; *p<0.05, **p<0.001, t-test). (B) Western blot analysis of Glut-1, Glut-5, Glut-2, Glut-7, Glut-9, and Glut-11 expression in LNCaP and PC-3 cells. Actin was used as a loading control. Molecular sizes (kDa) are indicated on the left side of the panel. (C) Densitometric quantification of Glut-1 and Glut-5 expression in LNCaP and PC-3 cells under glucose and fructose conditions (n=3-5; ns: not significant; *p<0.05, **p<0.001, t-test). (D) Immunofluorescence staining of Glut-1 and Glut-5 in LNCaP and PC-3 cells after 10 days of treatment with 5.5 mM glucose or 5.5 mM fructose. DAPI was used to counterstain nuclei. White arrows indicate intracellular Glut-5 expression. Scale bar: 10 µm. (E) Quantification of glucose transporter expression under glucose and fructose treatment, measured as integrated intensity normalized per cell (n=3; ***p<0.001, t-test). (F) Endpoint fluorescence measurement in LNCaP and PC-3 cells stimulated with 5.5 mM fructose versus 5.5 mM glucose for 10 days (n=3; **p < 0.0059, ***p < 0.0009).

To confirm our main results, immunofluorescence against Glut-1 and Glut-5 was performed under chronic fructose and glucose (control) stimulation (Fig. 2D). The abundance of Glut-5 was significantly increased in the fructose condition when compared to glucose in both cancer cell types, whereas Glut-1 protein was not affected (Fig. 2D-E). Interestingly, the expression of Glut-5 was not increased at the plasma membrane in both cells after fructose exposure. Therefore, we explored how fructose stimulation impacts the Glut-5 and EEA1 co-localization in LNCaP and PC-3 cells (Supp. Fig. 3). The Glut-5-EEA1 colocalization significantly decreased under fructose treatment in both LNCaP and PC-3 cell lines (Supp. Fig. 3B), which correlated with a significant increase in the expression of EEA1 (Supp. Fig. 3C) under fructose treatment. These data suggest that chronic fructose stimulation increases Glut-5 expression; however, the transporter remains largely intracellular, potentially indicating a defect or regulatory mechanism preventing its trafficking to the plasma membrane. We tested the fructose transport capacity of cells using a fluorescently-labelled ManCou-1 probe, which is specifically transported by Glut-5^29^. Our results demonstrated that chronic fructose exposure significantly increased ManCou-1 uptake in both LNCaP and PC-3 cells compared to glucose-treated controls (Fig. 1F). These findings suggest that fructose stimulation enhances SLC2A5/Glut5 expression but does not significantly modify its subcellular localization. This implies a potential role for Glut5 not only in fructose uptake but also in modulating intracellular signaling pathways. Alternatively, it remains possible that fructose-induced signaling activates a separate mechanism responsible for substantial mobilization of Glut5 to the plasma membrane in prostate cancer cells.

### Chronic fructose stimulation diminishes lactate and mitochondrial ATP production in prostate cancer cells

To expand our findings on the chronic effects of fructose on PCa cell metabolism, we examined the basic metabolic parameters, lactate and ATP production, in LNCaP and PC-3 cells. Through an empiric observation of our fructose treatments, we noticed that around 24-48 hours of treatment the phenol red from the cell medium demonstrated, in an indirect way, less acidification levels (lactate or HCO_3_^−^ production) into the cell medium from fructose *vs.* glucose treatment in both cell types (Fig. 3A). Based on this observation, we measured extracellular and intracellular lactate levels under long-term fructose exposure. We observed that extracellular lactate was reduced by more than 15-fold in the fructose condition compared to the glucose condition in LNCaP cells, and by 4-fold in PC-3 cells (Fig. 3B). In line with these results, intracellular lactate levels were 5-fold and 3-fold lower in the fructose condition compared to glucose treatment in both LNCaP and PC-3 cell lines, respectively (Fig. 3C).

**Figure 3.**
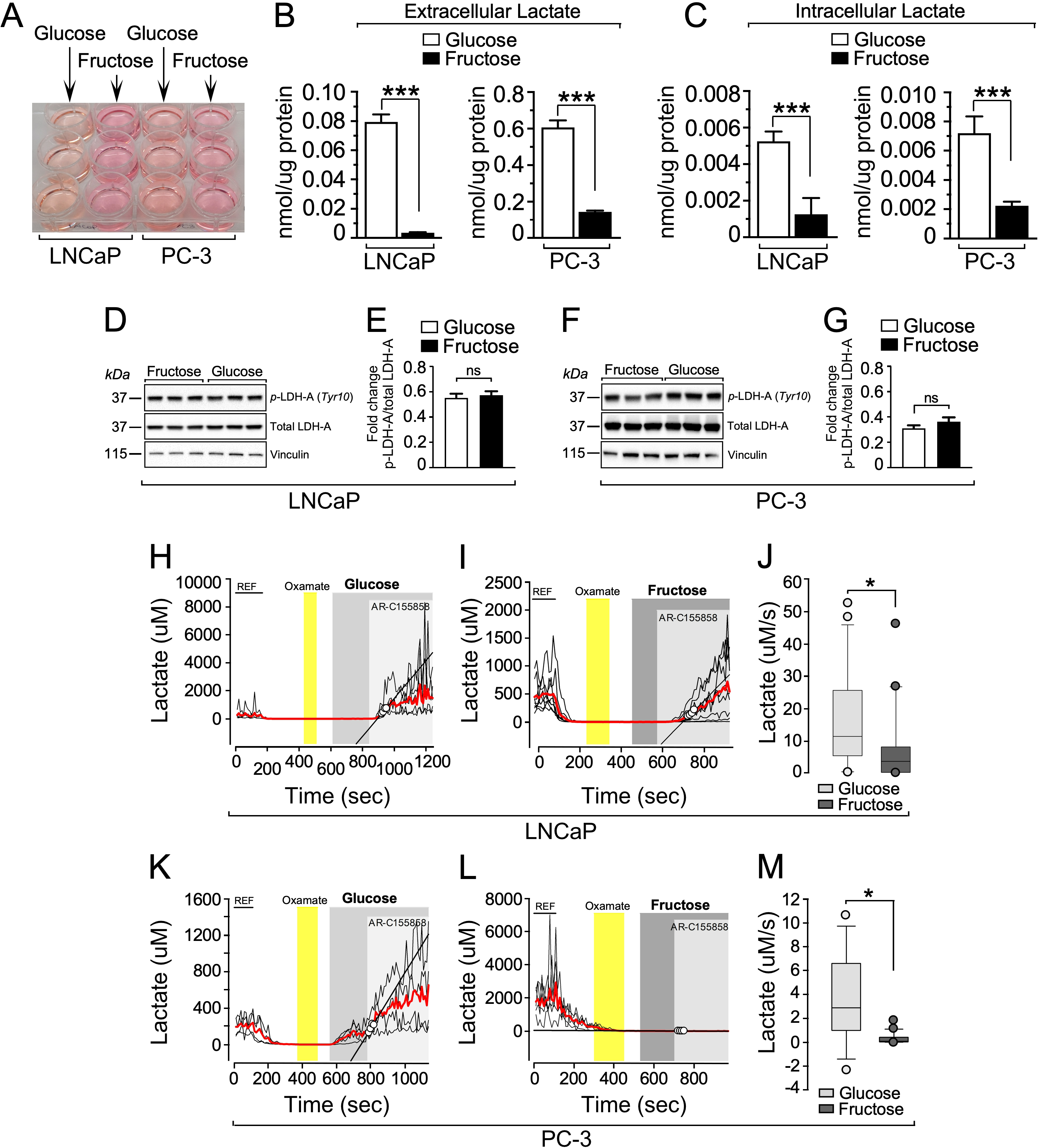
Chronic and acute fructose stimulation decreases lactate production in prostate cancer cells *in vitro*. (A) Representative images of cell culture plates showing LNCaP and PC-3 cells treated with 5.5 mM glucose or 5.5 mM fructose for 48 hours. (B) Extracellular lactate levels in LNCaP and PC-3 cells following 48 hours of treatment with 5.5 mM glucose (white bars) or 5.5 mM fructose (black bars) (n=3; ***p<0.001, t-test). (C) Intracellular lactate levels in LNCaP and PC-3 cells after 48 hours of treatment with 5.5 mM glucose (white bars) or 5.5 mM fructose (black bars) (n=3; ***p<0.001, t-test). (D) Representative western blot analysis of p-LDH-A (Tyr10) and total LDH-A expression in LNCaP cells treated with fructose or glucose for 48 hours. Vinculin was used as a loading control. Molecular sizes (kDa) are indicated on the left side of the panel. (E) Densitometric quantification of p-LDH-A (Tyr10) and total LDH-A protein expression in LNCaP cells under glucose (white bars) or fructose (black bars) conditions (n=3; ns: not significant, t-test). (F) Western blot analysis of p-LDH-A (Tyr10) and LDH-A expression in PC-3 cells treated with fructose or glucose for 48 hours. Vinculin was used as a loading control. Molecular sizes (kDa) are indicated on the left side of the panel. (G) Densitometric analysis of p-LDH-A (Tyr10) and LDH-A expression in PC-3 cells, showing no statistically significant difference between treatments (n=3; ns: not significant, t-test). (H) Intracellular lactate production flux experiment in LNCaP cells exposed to 1µM of MCT blocker AR-C155858 in the presence of 5 mM glucose or (I) 5 mM fructose. To calibrate the intracellular readout, lactate depletion was induced via trans-acceleration exchange by replacing the REF buffer (5 mM glucose and 1 mM lactate) with 6 mM oxamate. MCT: monocarboxylate transporters. (J) Summary data from n=3, with glucose tested in 24 cells and fructose in 21 cells (Mann-Whitney rank sum test *p = 0.016). (K) Intracellular lactate production flux experiment in PC-3 cells exposed to 1 µM of MCT blocker AR-C155858 in the presence of 5 mM glucose or (L) 5 mM fructose. To calibrate the intracellular readout, lactate depletion was induced via trans-acceleration exchange by replacing the REF buffer (5 mM glucose and 1 mM lactate) with 6 mM oxamate. (M) Summary data from n=3, with glucose tested in 12 cells and fructose in 20 cells (Mann-Whitney rank sum test *p <0.001).

To investigate whether the reduced lactate production in LNCaP and PC-3 cell lines was associated with a downregulation of lactate dehydrogenase (LDH) expression and/or activation, we measured total LDH protein levels and the phosphorylation status of LDH in these cells exposed to fructose using Western blot analyses (Fig. 3D and 3F). Our findings indicated that neither the total LDH levels nor the phosphorylation status of this enzyme were significantly altered following fructose treatment compared to glucose (Fig. 3E and 3G). These results suggest that exposure to fructose decreases lactate production in PCa cells, likely by redirecting metabolic flux toward pyruvate synthesis and thereby shifting the metabolic phenotype from glycolytic to oxidative.

Interestingly, our empirical observation of the culture medium suggests that changes in the intracellular production and release of lactate into the extracellular space, occur as early as 24 hours of incubation with fructose when compared to glucose. This prompted us to analyze the acute effect of fructose incubation on intracellular lactate production using Laconic, a genetically encoded fluorescent indicator for lactate^24^. Laconic enables the detection and quantification of lactate production flux with high spatiotemporal resolution^24^. We expressed Laconic in LNCaP and PC3 cells and exposed them to an acute pulse of glucose (Fig. 3H and 3K) or fructose (Fig. 3I and 3L). In presence of these carbon sources, the lactate production flux was assessed using a pharmacological stop-transport protocol with the monocarboxylate transporter inhibitor ARC-155858; this approach inhibits lactate export. Inhibition of lactate export perturbed the intracellular lactate steady-state, leading to lactate accumulation proportional to the rate of lactate production^30^. In both LNCaP and PC-3 cell lines, fructose significantly reduced the lactate production rate (Fig. 3J and 3M). Notably, lactate production in PC-3 cells dropped to nearly undetectable levels during acute fructose exposure (Fig. 3L and 3M). These findings suggest that fructose induced an acute rerouting of metabolic fluxes and a reversal of the Warburg phenotype in PCa cells.

Reduced glycolytic activity in the presence of fructose may be associated with increased pyruvate availability within the mitochondria and enhanced ATP production. To confirm mitochondrial activity in our experimental setup, we measured both extra- and intracellular ATP levels in LNCaP and PC-3 cells under basal conditions. The results revealed that PC-3 cells exhibited higher levels of both intracellular and extracellular ATP compared to LNCaP cells (Fig. 4A-B). Notably, chronic fructose stimulation led to a significant reduction in intracellular ATP levels in PC-3 cells, whereas no notable change was observed in LNCaP cells (Fig. 4C). Given that intracellular ATP levels reflect the cellular energy steady-state rather than direct ATP production, we proceeded to isolate mitochondria to assess their functionality by measuring ATP production and oxygen consumption (Fig. 4D). Our findings demonstrated that both mitochondrial ATP production (Fig. 4E) and oxygen consumption (Fig. 4F) were significantly reduced following chronic fructose stimulation. Collectively, these data suggest that fructose stimulation decreases both lactate and ATP production in LNCaP and PC-3 cells, potentially indicating that carbon flux is primarily directed towards *de novo* lipogenesis *via* citrate production derived from the Krebs cycle.

**Figure 4.**
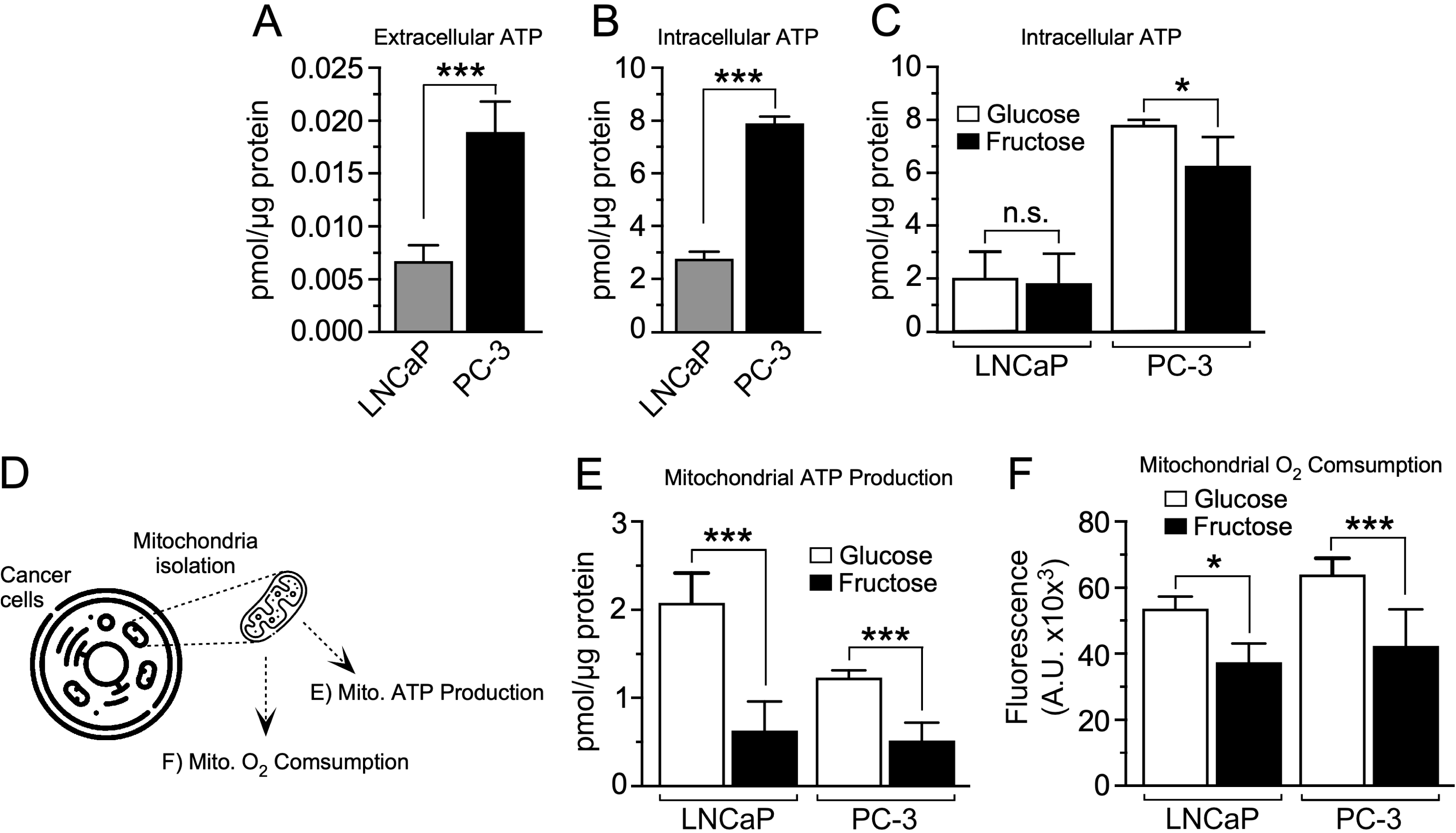
Fructose stimulation decreases ATP production and mitochondrial oxygen consumption in prostate cancer cells *in vitro.* (A) Extracellular and (B) intracellular ATP levels under basal conditions in LNCaP and PC-3 cells (n≥4; ***p<0.001, t-test). (C) Intracellular ATP levels following hexose stimulation for 10 days (n≥4; *p<0.05, t-test). (D) Graphic scheme of mitochondrial-enriched fraction isolation in prostate cancer cells. (E) Mitochondrial ATP production in LNCaP and PC-3 cells under glucose (white bars) and fructose (black bars) stimulation for 10 days (n=5; ***p<0.001, t-test). (F) Mitochondrial oxygen consumption in LNCaP and PC-3 cells under glucose (white bars) and fructose (black bars) stimulation for 10 days (n=3; *p<0.05, ***p<0.001, t-test).

### Chronic fructose stimulation enhances G6PD and FASN enzymes and promote lipid synthesis in prostate cancer cells

To elucidate the metabolic pathways altered by chronic fructose stimulation in PCa cells, we examined the expression levels of the metabolic enzymes FASN, TKT, and G6PD after 10 days of fructose stimulation in LNCaP and PC-3 cells. These enzymes are known to be key components of the anabolic pathways DNL and PPP^23,24,29^ (Fig. 5A). Notably, the expression of FASN and G6PD, but not TKT, transcripts exhibited an approximately 8.3-fold and 8.6-fold increase, respectively, under fructose stimulation compared to the control condition in LNCaP cells (Fig. 5B, LNCaP). In contrast, no statistically significant alterations were detected in the expression levels of these genes in PC-3 cells following fructose treatment (Fig. 5B, PC-3). These findings were corroborated through Western blot and immunofluorescence analyses. In the Western blot assays, FASN and G6PD exhibited a 2.2-fold and 6.0-fold increase, respectively, while TKT demonstrated a 0.5-fold decrease under fructose conditions compared to glucose in LNCaP cells (Fig. 5C-D, LNCaP). Notably, in PC-3 cells, fructose stimulation led to an increase in G6PD protein expression, with no significant changes observed in FASN or TKT (Fig. 5C-D, PC-3). Immunofluorescence analyses further supported these results, demonstrating that in LNCaP cells, FASN and G6PD were significantly upregulated, whereas TKT was downregulated in the presence of fructose (Fig. 5E-F, LNCaP). Conversely, in PC-3 cells, only G6PD showed a significant increase in expression under fructose conditions (Fig. 5E-F, PC-3). These findings suggest that fructose stimulation induces distinct enzymatic responses contingent upon the PCa cell line, potentially modulated by variations in their aggressiveness or androgen receptor status. Notably, the data indicate that the differential expression of FASN and G6PD under fructose stimulation may directly or indirectly contribute to the activation of the *de novo* lipogenesis (DNL) pathway.

**Figure 5.**
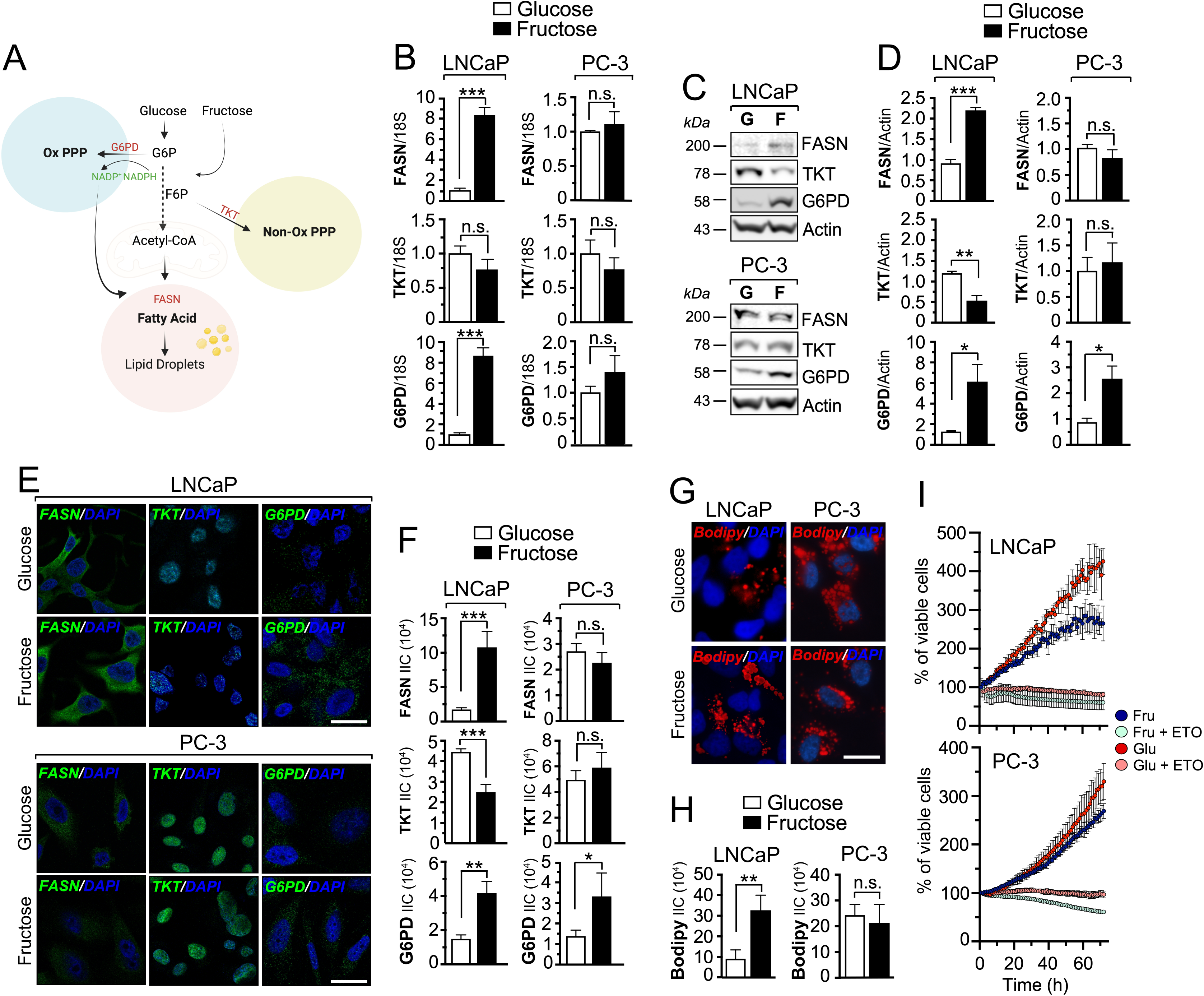
Chronic fructose stimulation enhances FASN and G6PD expression and promotes lipid droplet accumulation in LNCaP cells. (A) Schematic representation of key enzymes and metabolites involved in glycolysis, fructolysis, de novo lipogenesis (DNL), the non-oxidative pentose phosphate pathway (PPP), and the oxidative PPP. Red indicates key enzymes; green indicates electron donors. G6P: glucose-6-phosphate; F6P: fructose-6-phosphate; G6PD: glucose-6-phosphate dehydrogenase; TKT: transketolase; NADP: oxidized form of nicotinamide adenine dinucleotide phosphate; NADPH: reduced form of nicotinamide adenine dinucleotide phosphate; Acetyl-CoA: acetyl coenzyme A; FASN: fatty acid synthase. (B) mRNA levels of FASN, TKT, and G6PD in LNCaP and PC-3 cells following stimulation with 5.5 mM glucose or 5.5 mM fructose for 10 days (n=4-5; ns: not significant; ***p< 0.0005, t-test). (C) Western blot analysis of FASN, TKT, and G6PD expression in LNCaP and PC-3 cells. Actin was used as a loading control. Numbers on the left side of the panel indicate molecular weight (kDa). (D) Densitometric quantification of FASN, TKT, and G6PD protein levels in LNCaP and PC-3 cells under stimulation with 5.5 mM glucose or 5.5 mM fructose for 10 days (n=4-5; ns: not significant; *p<0.05; **p<0.001; ***p<0.0001, t-test). (E) Immunofluorescence staining of FASN, TKT, and G6PD in LNCaP and PC-3 cells treated with 5.5 mM glucose or 5.5 mM fructose for 10 days. DAPI was used to counterstain nuclei. White bar: 10 µm. (F) Integrated intensity, normalized per cell, quantifying FASN, TKT, and G6PD protein expression under 5.5 mM glucose or 5.5 mM fructose (n=3; ns: not significant; *p<0.05; **p<0.001; ***p<0.0001, t-test). (G) Representative images of ORO staining in LNCaP and PC-3 cells treated with 5.5 mM glucose or 5.5 mM fructose for 10 days. DAPI was used to counterstain nuclei. White bar: 10 µm. (H) Quantitative comparison of the staining effect by number of lipid droplet formation normalized per cell. (I) Effect of the lipid metabolism inhibitor Etomoxir (ETO) on fructose-stimulated or glucose-stimulated LNCaP and PC-3 cells.

Since FASN and G6PD are two enzymes closely associated with lipid homeostasis^31,32^, and were upregulated in the less aggressive, androgen-sensitive, LNCaP cells under fructose stimulation, we sought to further investigate this phenomenon by analyzing lipid droplet (LD) accumulation in both cell lines. LNCaP cells demonstrated a significant increase in the number of lipid droplets under fructose stimulation compared to glucose conditions (Fig. 5G, H, LNCaP). In contrast, lipid droplet accumulation in PC-3 cells remained unchanged across both experimental conditions. It is worth noting that LD accumulation in PC-3 cells was already elevated under control (glucose) conditions (Fig. 5G, PC-3). Collectively, these findings suggest that fructose stimulation, activates the DNL pathway and promotes LD accumulation in the less aggressive, androgen-sensitive, LNCaP cell line.

Our findings reveal that both cell lines, LNCaP under fructose stimulation and PC-3 under both basal and fructose-stimulated conditions, exhibit a marked capacity to accumulate significant lipid stores (Fig. 5G-H). This observation underscores a strong reliance on lipid metabolism in both cell types. To substantiate this hypothesis, we evaluated the proliferative capacity of both cell lines in the presence of the lipid metabolism inhibitor etomoxir under fructose-stimulated conditions (Fig. 5I). Our results demonstrated that etomoxir completely inhibited the proliferation of both LNCaP and PC-3 cell lines (Fig. 5I). These findings highlight the critical role of lipid metabolism in PCa cells, emphasizing its dependency under both basal and fructose-stimulated conditions, which suggests that targeting lipid metabolism could represent a promising therapeutic strategy to curb PCa progression.

Lastly, to determinate the effects of fructose on the metabolite profile of PCa cell lines, we carried out metabolomic analysis by using the Biocreates pipeline, following chronic glucose or fructose exposure in LNCaP and PC-3 cells. In figure 6A, different metabolites related to amino acid such as ceramides, arginine, serine, and alanine were highly abundant in LNCaP cells versus PC-3 cells independent of the hexose treatment. On the other hand, metabolites such as tryptophan, histidine and γ-aminobutyric acid (GABA) were highly abundant in the aggressive PC-3 cell line and lower abundancy in the androgen dependent cell line, LNCaP. This data suggests that LNCaP and PC-3 cell lines rely on different metabolic pathways, been amino acid and GABA, the main metabolic pathways, respectively. Moreover, we were able to detect three major metabolites highly abundant in LNCaP cells treated with fructose, triglyceryl palmitate (TG), eicosapentaenoic acid (EPA) and docosahexaenoic acid (DHA), when compared to the glucose group (Fig 6B). In the case of PC-3 cells under fructose treatment, metabolites such as 5-aminovaleric acid (AVA), ceramide 5, ceramide 18 and valine were highly abundant in relation to the glucose group. Interestingly, in the glucose group, PC-3 cells presented already high levels of 1,2-dipalmitoyl-glycerol (DG: 16:0, DG: 16:1), eicosatrienoic acid (FA: 20:3) and homocysteine (HCys) (Fig 6C). Together this data suggest that fructose stimulation enhances lipid metabolism by either upregulating key enzymes of the DNL pathway which can be reflected with key metabolites products such as TG, EPA and DHA in LNCaP cells, where fructose stimulation in PC-3 cells, suggests not to affect the generation of new lipid metabolites.

**Figure 6.**
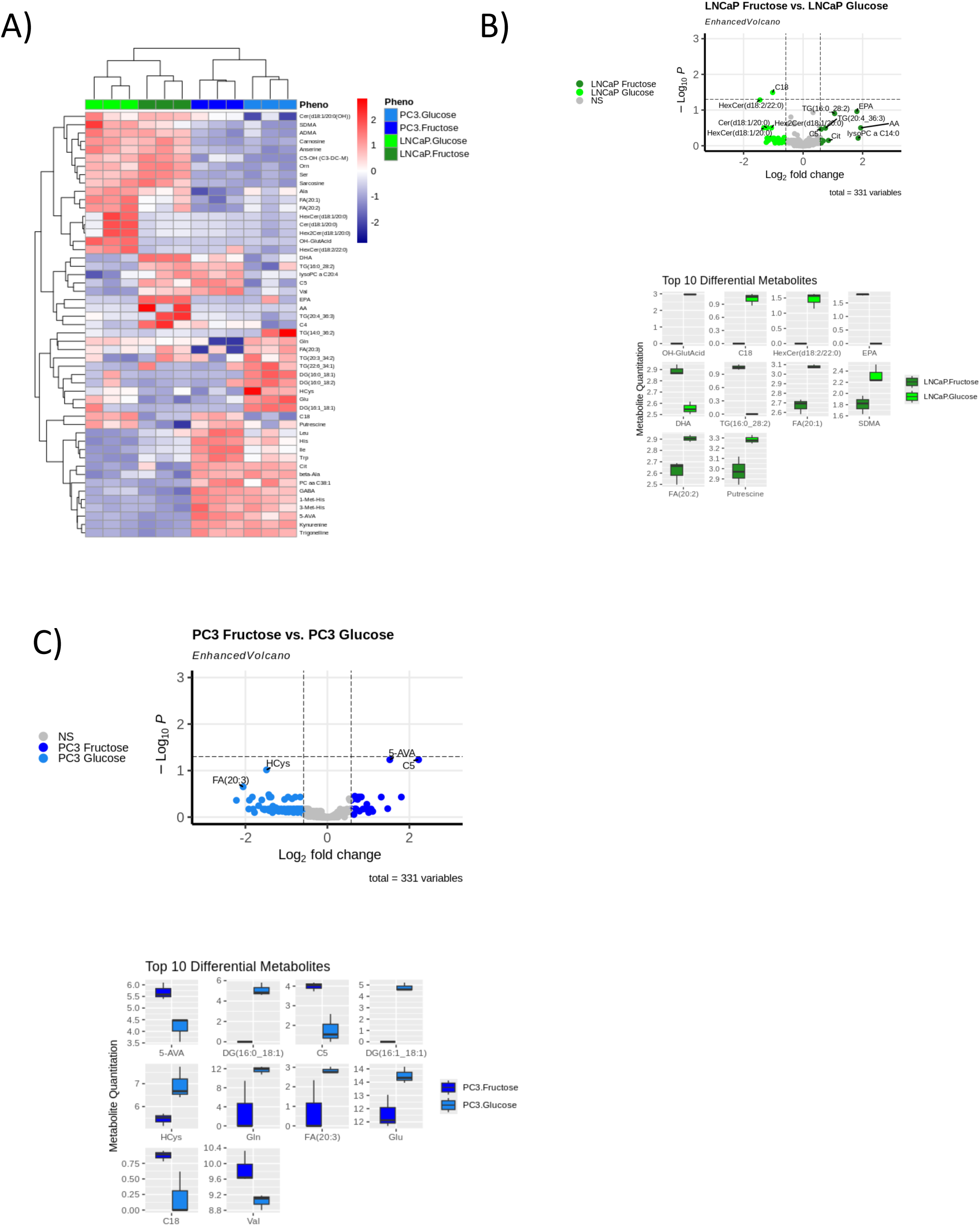
Fructose stimulation enhances lipid metabolism in LNCaP cells under fructose treatment. (A) Heatmap of significantly altered metabolites in LNCaP and PC-3 cells under different hexose stimulations. The colors in the heat map indicates the log transformed values of each metabolite. (B) Volcano plots depicting metabolomic diversity in each hexose stimulation. Each point represents a metabolite. Decreased (log2(FC)≤ −1.0, left side of dotted vertical lines) or increased (log2(FC)≥1.0, right side of dotted vertical lines. Box plots with the top 10 differential metabolites in LNCaP cells under glucose (light green) versus fructose (dark green) stimulation for 10 days. (C) Volcano plots and box plot with the top 10 differential metabolites in PC-3 cells under glucose (light blue) or fructose (dark blue) stimulation for 10 days. Medians are indicated by horizontal lines withing each box n=3. NS: no significance.

## Discussion

A growing body of research explores fructose’s effects on cancer, from molecular mechanisms to systemic impacts in animal models^6,11,12,21^. While metabolic alterations are key in cancer progression, few studies have systematically examined fructose’s effects on cancer metabolism, and none on PCa cells. To address this gap, we conducted an in vitro study using two PCa cell lines: LNCaP (low-aggressiveness) and PC-3 (high-aggressiveness). This approach allowed us to assess distinct metabolic responses to fructose and its potential role in PCa progression.

Under basal conditions, our immunofluorescence coupled with confocal analysis revealed that Glut-5 was predominantly localized at the cytoplasmic level in both LNCaP and PC-3 cells. This observation was further substantiated by the colocalization of Glut-5 with transferrin and, more specifically, with the early endosomal marker EEA1^33^. In support of this observation, previous studies from our group^34–36^ and others^37–39^ have consistently reported that Glut-5 protein in malignant cells from human tissue specimens of various origins is predominantly localized in the cytoplasm. Moreover, immunogold analyses reported by Godoy *et al*.,^19^ in breast cancer tissue demonstrated that Glut-5 was localized intracellularly within vesicle-like structures in malignant cells and in breast cancer cell lines. Notably, despite this accumulated evidence, no studies have sought to clarify the specific localization of the so-called “cytoplasmic immunostaining”, typically attributed to Glut-5, in cancer cells. Our study, for the first time, indicates that this cytoplasmic immunostaining may reflect a localization of Glut-5 within the early endosomal compartment, an intracellular membranous structure that plays a critical role in the turnover of membrane receptors and transporters^40,41^. These findings raise several questions regarding the biology of this transporter. For example, does its localization within early endosomal compartments indicate a slower trafficking process toward its final destination at the plasma membrane? Alternatively, and perhaps more intriguingly, could this protein undergo recycling from the plasma membrane and be stored intracellularly, only to be redeployed to the plasma membrane in response to specific stimuli or external conditions? Although definitive evidence supporting either hypothesis is currently lacking, previous studies in benign mouse enterocytes suggest that this transporter may translocate from intracellular storage compartments to the apical membrane in response to its substrate, fructose^37^. This process involves Ras-related protein-in-brain 11 (Rab11)-dependent endosomes and requires the presence of both Glut-5 and KHK, indicating that fructose transport and its intracellular metabolism are necessary to modulate Glut-5 localization.

Based on our findings, it is reasonable to hypothesize that cancer cells may exhibit a similar response under fructose stimulation. Liang *et al.*^12^ found that VAMP8 (vesicle associated membrane protein 8) was upregulated in fructose trained PC-3 cells (after > 20 passages). VAMP8 is identified as an endosomal protein that participates in diverse biological functions including endosomal fusion, granule-to-granule fusion and autophagy^42^. Although our data did not confirm potential translocation of Glut-5 from endosomal compartment to the plasma membrane under chronic fructose stimulation, our results could be limited by the duration of fructose exposure. Further studies employing advanced microscopy techniques or live-cell imaging analyses will be required to elucidate this highly dynamic cellular process.

Another plausible explanation for the preferential localization of Glut-5 in the endosomal compartment is its potential role as a transceptor in PCa cells under fructose stimulation. Transceptors function as both transporters and signaling receptors, where substrate binding—in this case, fructose—not only facilitates transport but also activates intracellular signaling pathways. This signaling triggers adaptive responses that align the expression of transporters and metabolic enzymes with nutrient availability essential for cancer cell growth and survival^43–45^. The transceptor role of Glut-5 has been indirectly demonstrated in *ex vivo* intestinal organoids^46^. In this model, fructose introduced via the basolateral membrane failed to stimulate fructose-responsive genes in organoids derived from Glut-5^−/-^ mice, suggesting that Glut-5 is involved in fructose sensing, even though it is not required for basolateral fructose transport into the cytosol. Understanding the role of transceptors in fructose metabolism will provide insight into cancer metabolism and potential therapeutic targets.

In our study, fructose but not glucose increased the expression of Glut-5 in LNCaP and PC-3 cells, underscoring the specificity of fructose in driving this effect. These findings align with the study by Liang et al.^12^, which reported a robust upregulation of Glut-5 expression in PC-3 cells chronically stimulated with fructose over 10 cell passages. Notably, our results revealed that the expression levels of the glucose transporter Glut-1, as well as other fructose transporters (Glut-2, Glut-7, Glut-9, and Glut-11), remained unchanged despite chronic fructose stimulation, highlighting the specificity of fructose in inducing Glut-5 expression. Despite these intriguing observations, the molecular mechanisms by which fructose specifically induces the upregulation of Glut-5 in PCa cells remain largely unknown. Given the specificity of *SLC2A5* overexpression, it is plausible that extrinsic or epigenetic factors affecting the *SLC2A5* locus contribute to this upregulation. For instance, inflammatory factors such as IL-6 have been shown to activate fructose uptake via STAT3, and its interaction with the Glut-5 promoter region has been demonstrated to enhance Glut-5 transcription in oral squamous cell carcinoma^47^. Another key element to add regarding regulation of Glut-5 expression, is hypoxia. The regulation of Glut-5 by Hif1-α is not limited to specific cancer types, but represent a clear phenomenon of a metabolic adaptation to hypoxia^48^. This regulatory mechanism permits cancerous cells to increase fructose uptake, supporting survival under low oxygen, and probably low glucose, conditions^49,50^. The prostate gland is not inherently hypoxic under normal physiological conditions^14^; however, the prostate can become hypoxic and develop inflammation in certain situation such as PCa or benign prostatic hyperplasia (BPH). Understanding inflammation and hypoxic pathways, and the effect of fructose under these conditions, are key areas of research in the male reproductive tract, especially considering that an increase in the consumption of dietary sugar, particularly in the form of HFCS^4,51–53^ have demonstrated a direct relationship with a higher risk of developing PCa or symptomatic PCa.

Carreño *et al*.^10^ demonstrated that downregulation of Glut-5 expression using siRNA consistently attenuated the effect of chronic fructose exposure on cell proliferation in PCa cells. This finding suggests that, at least in part, the pro-proliferative effects of fructose in PCa cells rely on intracellular mechanisms that involve changes in energy metabolism^54,55^. Consequently, we analyzed the impact of fructose on key metabolic outputs, specifically focusing on lactate production and ATP levels. Our study revealed that fructose exposure significantly modifies the metabolic phenotype of PCa cells, notably by reducing intra- and extracellular lactate levels, decreasing ATP production, and potentially redirecting carbon flux toward lipogenic pathways, with potential implications for PCa cell survival and proliferation^12^. It has been described that reduction of total hexose and lactate accounts in the liver indicates impaired glycolytic flux^56^, which can contribute to the development of chronic inflammation and trigger different pathologies such as nonalcoholic steatohepatitis^57^, thus having lower levels of lactate in the presence of fructose in our PCa cells, strongly suggest that fructose is been used to lipid formation.

Chronic fructose stimulation was associated with reduced intracellular ATP levels in PC-3 cells but not in LNCaP cells, highlighting cell line-specific differences in metabolic responses. However, these levels reflected the steady-state ATP production characteristic of each cell line and did not specifically represent mitochondrial ATP production^58^. Consequently, we measured the ATP production on isolated mitochondrion of fructose-stimulated LNCaP and PC-3 cells. We found that chronic fructose stimulation diminished mitochondrial activity, as evidenced by the decreased mitochondrial ATP production and oxygen consumption observed in fructose-treated cells, potentially redirecting pyruvate-derived carbons toward lipogenesis via citrate intermediates^59^. Decreased ATP production after fructose stimulation has been observed in other cell models. Latta and colleagues^60^ observed ATP depletion using primary hepatocytes by different phosphates-trapping carbohydrates, including fructose. Furthermore, pronounced ATP depletion (75-80%) was also observed after addition of carbohydrates metabolized via aldolase pathway, such as sorbitol, tagatose and 2,5-anhydro-mannitol, whereas no ATP depletion was observed after addition of glucose, or mannose, which are hardly metabolized via the aldose pathway.

The decrease in ATP production may reflex a potential shift in carbon flux, toward de novo lipogenesis, which highlight the metabolic flexibility of PCa cells in adapting to nutrient availability^61^. Citrate, a key intermediate of the Krebs cycle is often utilized for lipid biosynthesis in PCa cells to support membrane formation and energy storage. By enhancing citrate production through oxidative metabolism, fructose may promote lipogenesis at the expense of ATP synthesis, which could influence tumor growth and aggressiveness^62^. The ability of fructose to modulate metabolic pathways in PCa cells underscores its potential role in influencing tumor progression.

We demonstrated that chronic fructose stimulation increases the expression of FASN and G6PD enzymes at both the mRNA and protein levels, while a decrease in TKT expression was observed in LNCaP cells. Interestingly, in PC-3 cells, only the G6PD enzyme was upregulated under fructose stimulation. These findings highlight two important aspects of PCa biology: 1) fructose may promote de novo lipogenesis (DNL) in PCa cells, and 2) fructose stimulation differentially impacts PCa cell lines, likely depending on their aggressiveness and/or androgen receptor (AR) signaling status.

FASN is responsible for the synthesis of palmitate necessary to fuel DNL pathway^6,63^. Different studies have linked fructose consumption with the generation of high-grade non-alcoholic fatty liver, through a mechanism that involves DNL stimulation through a steroid regulatory element binding protein (SREBP), which consequently can increase the level of several enzymes, including FASN^64,65^. In the context of prostate malignancy, FASN is described as a bona fide oncogene which is overexpressed in prostate intraepithelial neoplasia and metastatic PCa^66^. In addition to this, overexpression of FASN together with expression of the AR in prostate epithelial cells facilitates the development of invasive cancers when these cells are injected orthotopically into mice^67^. Interestingly, inhibition of FASN gene expression in various cancer cell lines via either RNA interference-mediates silencing or chemical inhibition induces intrinsic apoptosis, suggesting that FASN overexpression may protect prostate epithelial cells from apoptosis^68,69^. Our results suggest that under fructose stimulation, increased FASN expression was observed only in the androgen dependent cell line LNCaP but not in PC-3 cells, which may reflect a dependency on the AR signaling pathway, enhancing the DNL pathway by the increase of LD in LNCaP cells and potentially offering a protection by inhibiting the pathway of apoptosis.

It has been described that AR signaling in PCa can promote the accumulation of LDs^70^ and expression of enzymes such as FASN^71^. However, no study has delved into the potential relationship between fructose and AR signaling. AR determines bioenergetic traits through regulation of component in the glycolytic pathway (Glut-1, hexokinase I/II) and pyruvate flux into mitochondria (PDH). As an example, AR signaling increases expression of G6PD which directs glucose-6-phosohate from glycolysis to the PPP for generation of NADPH^72,73^. Thus, AR is capable to coordinately regulate the energy production and biosynthesis at multiple levels, suggesting that these metabolic pathways can act as potential targets to inhibit growth of PCa cells. Interestingly, a lack of effect of fructose on activating expression of DNL enzymes in PC-3 cells could be related to the lack of expression of AR. AR is capable to regulate fatty acid (FA) metabolism by controlling expression for more than 20 enzymes involves in many aspects of lipid metabolism, from uptake to degradation^67,74^. AR and the sterol regulatory-element binding protein (SREBP) regulate each other in a positive feedback system^73,74^. SREBP’s can target ELOV6 and SCD1, while FASN is targeted by both SREBP and AR. Thus, AR activation and fructose stimulation can accelerate fatty acid synthesis, particularly as the form of monounsaturated and saturated FA, such TG, EPA and DHA in LNCaP cells, which can be used as phospholipids, signaling molecules and storage molecules. Nevertheless, pharmacological or genetic inhibition of AR or enzymes related to the DNL needs to be done in order to demonstrate that these two pathways, AR and fructolysis, feed each other.

Overexpression of glucose-6-phosphate dehydrogenase (G6PD) enhances tumor cell resistance to cell death, primarily by regenerating NADPH through the oxidative branch of the PPP pathway, which is widely recognized as a cancer-promoting mechanism^75^. Remarkably, our results showed that chronic stimulation with fructose upregulate G6PD expression in LNCaP and PC-3 cell lines, suggesting a potential role of fructose in driving metabolic reprogramming toward enhanced oxidative PPP activity. This fructose-induced upregulation of G6PD may likely contribute to the maintenance of redox homeostasis, increased biosynthetic capacity, and tumor cell survival. Studies have shown that disruption of G6PD activity through inhibitors such as dehydroepiandrosterone (DHEA) leads to dysregulation of de novo lipogenesis (DNL), accompanied by reduced migration and proliferation capabilities in HeLa cells^75,76^. Notably, the polymerization of acetyl-CoA molecules into fatty acid chains during DNL relies heavily on NADPH derived from the oxidative PPP, as observed in metabolic processes within brown adipose tissue^77^. These findings highlight the integral role of G6PD in lipid biosynthesis and its broader implications in cellular metabolism and cancer biology. Collectively, these findings suggest that fructose can stimulate de novo lipogenesis (DNL), through the overexpression of two key enzymes, FASN and G6PD. These enzymes, functionally located in different metabolic pathways, contribute to LD accumulation in LNCaP cells and it might be dependent of the AR pathway.

Polyunsaturated fatty acids, such as DHA and EPA are increased in plasma lipids and blood cell membranes in response to fatty acids supplementation over the course of 12 months^78^, thus seen the increase of these metabolites in the LNCaP cells under the fructose treatment, suggest that fructose acts as a booster of lipid metabolic dependencies in an AR-dependent manner. On the other hand, PC-3 cells presented higher levels of different ceramides, which represents a heterogenous group of lipids that are identified by the specific fatty acyl moiety bonded to sphingosine with an amide bond^79^. Ceramides, can act as secondary messengers for different cellular signaling pathways, proliferation, migration and mitochondrial regulation^80^. Further studies are clearly indicated to define the interrelationships between fructose, FAs, and steroid hormones in prostate cancer development and progression.

In summary, we have demonstrated how prostate cancer (PCa) cell metabolism undergoes significant changes under both acute and chronic fructose stimulation. Notably, fructose treatment increases Glut-5 expression, although not its cellular localization, suggesting that Glut-5 may function as a transceptor. Additionally, our findings strongly indicate that androgen receptor (AR) signaling and fructose stimulation drive distinct alterations in fatty acid metabolism, including disruptions to the de novo lipogenesis (DNL) pathway. This raises the critical question of what directs AR toward specific metabolic preferences. Potential underlying mechanisms include the interplay of fructose, Glut-5, AR activity, differential actions of AR co-regulators, tumor microenvironment factors, and epigenetic modifications. For instance, a lipid-rich tumor microenvironment could promote cancer immune tolerance, as immunosuppressive immune cell subtypes—such as regulatory T cells and M2 macrophages—rely on elevated fatty acid oxidation, potentially using fructose as a signaling molecule^45^. Understanding and targeting the fructolysis pathway and/or the AR-metabolome axis presents a unique therapeutic opportunity, particularly for AR-driven, castration-sensitive PCa.

## Competing interest

The authors declare no conflict of interest.

## Acknowledgments

We thank Dr. Marcus D. Goncalves for critical reading of the article. This study was supported by Fondecyt 1221067 to A.G. C.E.E was supported by a PhD national fellowship from ANID-Chile N° 21210701.

---

**Supplementary Figure 1. Validation of Glut-5 antibody specificity.** (A) Glut-5 immunofluorescence was performed in the presence of a Glut-5 blocking peptide (BP) at two different concentrations (100 and 200 µg/mL). An Alexa-555-conjugated secondary antibody (red) was used to visualize Glut-5 expression in LNCaP and PC-3 cells. DAPI (blue) was used to stain nuclei. The absence of the primary antibody served as a negative control (Control). (B) Quantification analysis in LNCaP and PC-3 cells was conducted based on the integrated fluorescence intensity of each image, normalized to cell number and experimental condition. (C) Glut-1 immunofluorescence was performed in the presence of BP (200 µg/mL) in LNCaP and PC-3 cells. (D) Quantification analysis in LNCaP and PC-3 cells was conducted based on integrated fluorescence intensity. Bars represent mean ± SEM of three independent experiments (ns: not significant, t-test). White bar: 20 µm.

**Supplementary Figure 2. Effects of fructose on prostate cancer cell viability.** Cell viability, represented by the percentage of live cells (white bars) and dead cells (black bars), was assessed using the LUNA automated cell counter in LNCaP and PC-3 cells exposed to 5.5 mM glucose or 5.5 mM fructose at 0, 24, 48, and 96 hours. For each condition, both adherent and suspended cells were collected and analyzed.

**Supplementary Figure 3. Fructose enhances EEA1 expression in prostate cancer cells.** (A) Immunofluorescence staining of Glut-5 and EEA1 in LNCaP and PC-3 cells following 10 days of treatment with 5.5 mM glucose or 5.5 mM fructose. DAPI was used to counterstain nuclei. White scale bar: 10 µM. (B) Quantification of Glut-5 and EEA1 colocalization in LNCaP and PC-3 cells under glucose or fructose treatment (n=3; **p<0.01; one-way ANOVA, Manders coefficient). (C) Integrated intensity analysis of EEA1 expression, normalized per cell, under glucose and fructose treatment (n=3; **ns: not significant; **p<0.001; ***p<0.0001, t-test).

**Supplementary Table I:**
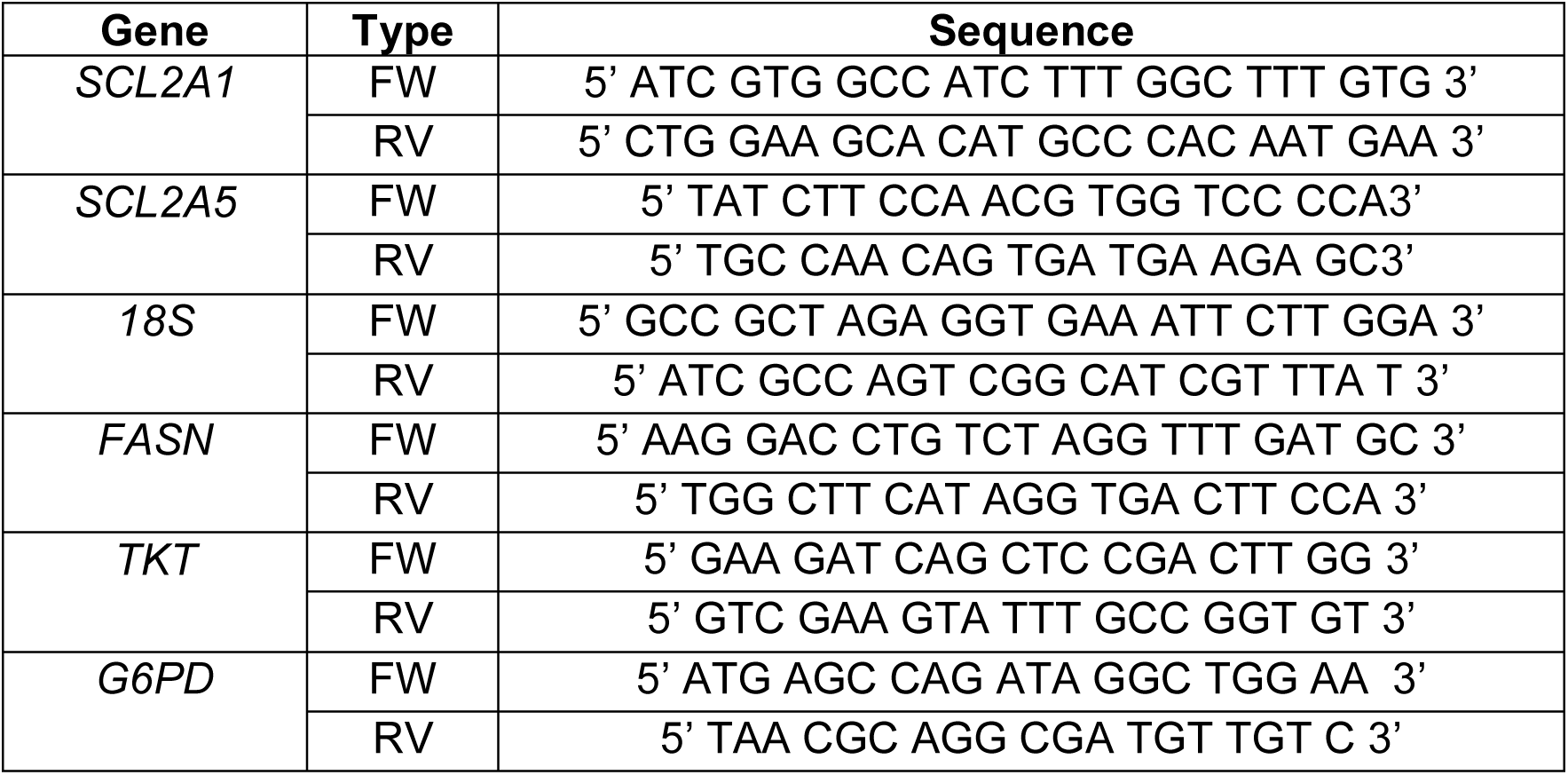

## Notes

### Competing Interest Statement

The authors have declared no competing interest.

