## Supplementary figures and images for "Metabolic Adaptations of Prostate Cancer Cells Under Chronic Fructose Stimulation"

### Supplemental FIgure 1

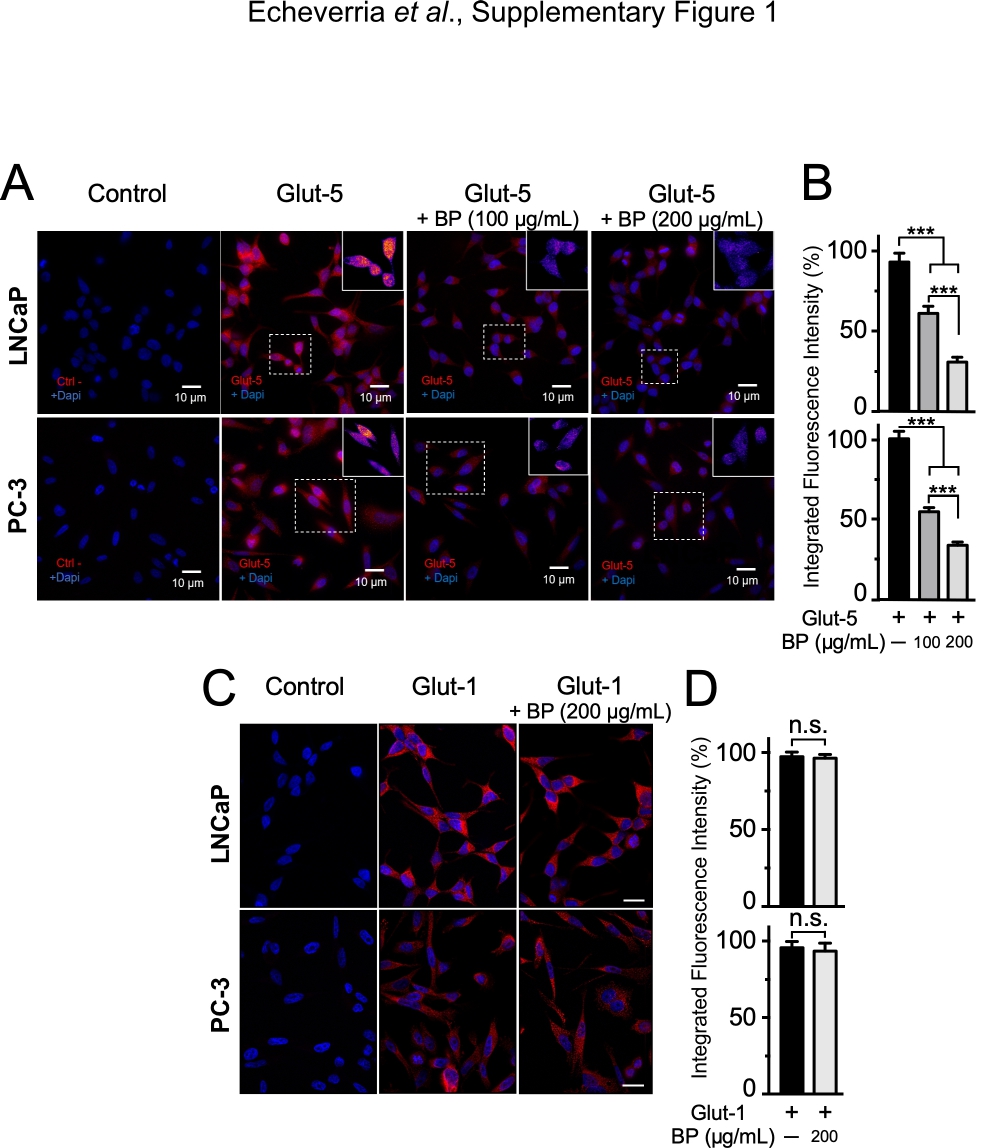

### Supplemental FIgure 2

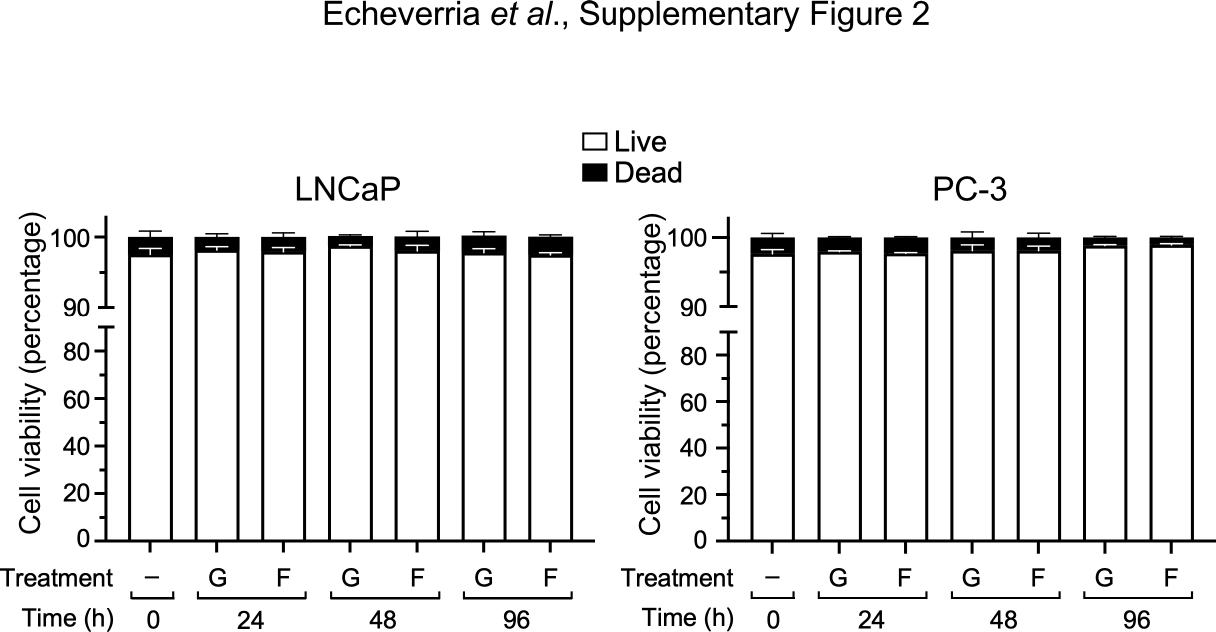

### Supplemental FIgure 3

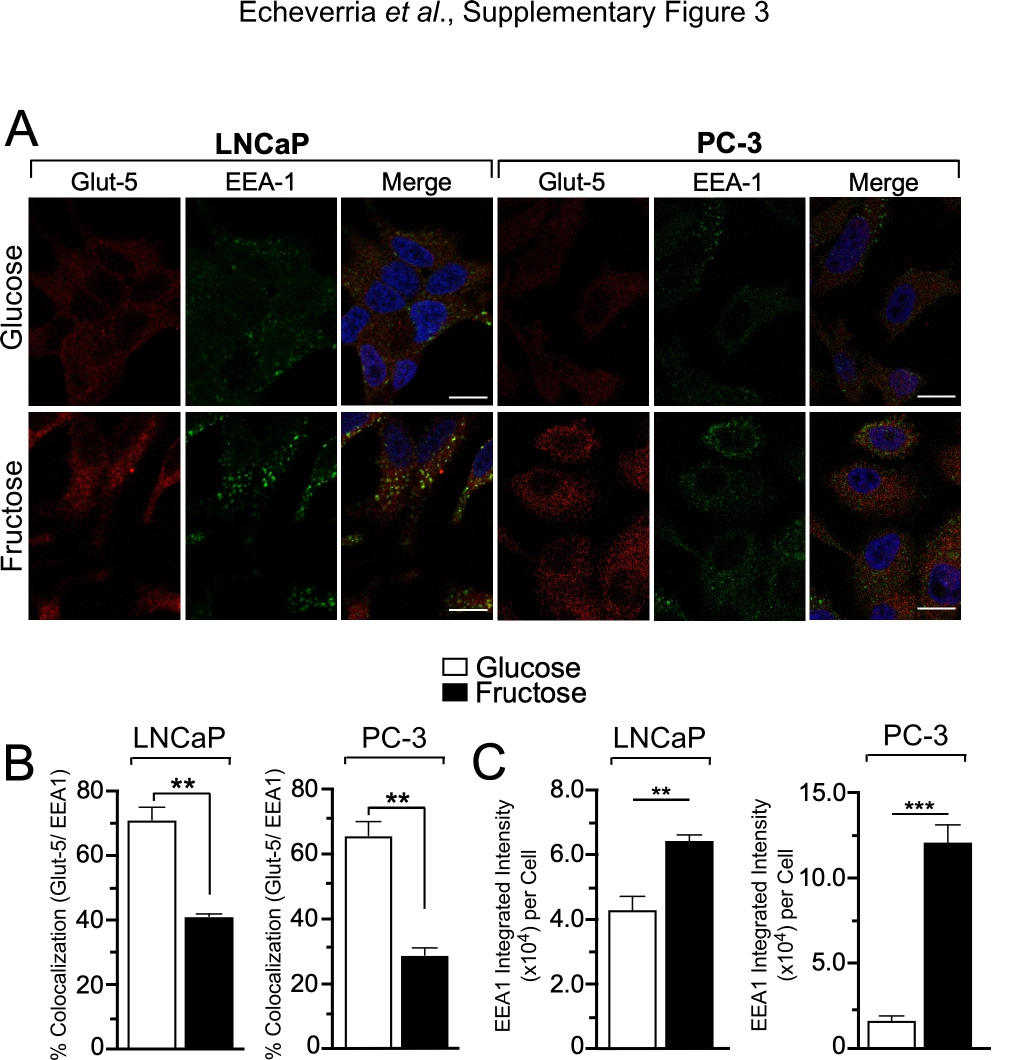
